# Disease-related single-point mutations alter the global dynamics of a tetratricopeptide (TPR) a-solenoid domain

**DOI:** 10.1101/598672

**Authors:** Salomé Llabrés, Ulrich Zachariae

## Abstract

Tetratricopeptide repeat (TPR) proteins belong to the class of α-solenoid proteins, in which repetitive units of α-helical hairpin motifs stack to form superhelical, often highly flexible structures. TPR domains occur in a wide variety of proteins, and perform key functional roles including protein folding, protein trafficking, cell cycle control and post-translational modification. Here, we look at the TPR domain of the enzyme O-linked GlcNAc-transferase (OGT), which catalyses O-GlcNAcylation of a broad range of substrate proteins. A number of single-point mutations in the TPR domain of human OGT have been associated with the disease Intellectual Disability (ID). By extended steered and equilibrium atomistic simulations, we show that the OGT-TPR domain acts as a reversibly elastic nanospring, and that each of the ID-related local mutations substantially affect the global dynamics of the TPR domain. Since the nanospring character of the OGT-TPR domain is key to its function in binding and releasing OGT substrates, these changes of its biomechanics likely lead to defective substrate interaction. Our findings may not only help to explain the ID phenotype of the mutants, but also aid the design of TPR proteins with tailored biomechanical properties.

## INTRODUCTION

Solenoid proteins represent ~5% of the human proteome. Due to their extended water-exposed surface and high degree of flexibility, they play a particularly important role in the formation of multiple protein-protein binding interactions. α-solenoid domains consist of arrays of repetitive α-helical units with variations in the numbers of repeats and the precise spatial arrangement of the helices [1]. Globally, most α-solenoid domains adopt extended superhelical shapes. The most common types of repetitive α-helical units are tetratricopeptide (TPR), HEAT, armadillo and leucine-rich repeats. Each repeat type possesses a characteristic conserved sequence of amino acids, which determines the specific fold of the units and influences the geometry and dynamics of the entire domain [2].

In the case of HEAT repeat domains, simulations have previously shown that the conserved hydrophobic core formed by part of this consensus sequence confers fully reversible, spring-like elasticity to the domains [3]. Armadillo repeat domains, by comparison, are more rigid, although they are still sufficiently flexible to accommodate a range of different binding partners [4]. The computationally predicted nanospring behaviour of alpha-solenoid domains has been experimentally confirmed for a designed protein consisting of three TPR repeats [5]. Altogether, it appears likely that the consensus sequence of the repetitive units governs the global dynamics of the domain. Depending on the specific functional role of the domain, this enables fine-tuning of its flexibility, while retaining stability against unfolding [5,6].

The TPR consensus sequence contains 34 residues, in which the conserved positions are W_4_-L_7_-G_8_-Y_11_-A_20_-F_24_-A_27_-P_32_ [7,8]. The sequence folds into a characteristic α-helix-turn-α-helix (helices A and B) motif [9]. TPR domain proteins are involved in a wide range of cellular processes such as protein folding [9–12], cell cycle control [13], post-transcriptional modification [14] and mitochondrial and peroxisomal protein transport [15,16]. Evolutionarily, the domains are likely to have arisen from the amplification of an ancestral helical hairpin structure [17]. Next to naturally occurring TPR domains, engineered TPR proteins have recently gained substantial interest as they allow the design of optimised protein-protein assembly surfaces [18,19].

O-linked GlcNAc-transferase (OGT; Fig. 1A) is a TPR-domain containing enzyme that catalyses O-GlcNAcylation, a reversible post-transcriptional modification of important protein substrates including transcription factors and cytoskeletal proteins [20]. The OGT-TPR domain recognises and binds substrate proteins and must therefore be able to adapt to a wide range of different protein sizes and geometries [21]. This capacity is shared with other α-solenoid domains, such as the HEAT and armadillo repeat domains that bind cargo proteins as nuclear transport receptors [22,23]. The OGT-TPR domain possesses an extended consensus sequence (N_6_-L_7_-G_8_-G_15_-A_20_-Y_24_-A_27_-Ψ_30_-P_32_), which includes three additional positions compared to most other TPR repeats [24] (Fig. 1B). Three single point mutations that are associated with Intellectual Disability (ID) phenotypes are located within repeat units TPR7 (L254F), TPR8 (R284P) and TPR9 (A319T), far from the catalytic domain of the enzyme [25] (Fig. 1A,C). Intellectual disability is a disease which leads to an early-onset impairment of cognitive function and the limitation of adaptive behaviour [26]. The X-ray structures of both the wild-type protein (wt, PDB id: 1W3B) [21] and the ID-associated OGT mutant L254F (PDB ID 6EOU) [14], have recently been determined.

**Figure 1.**
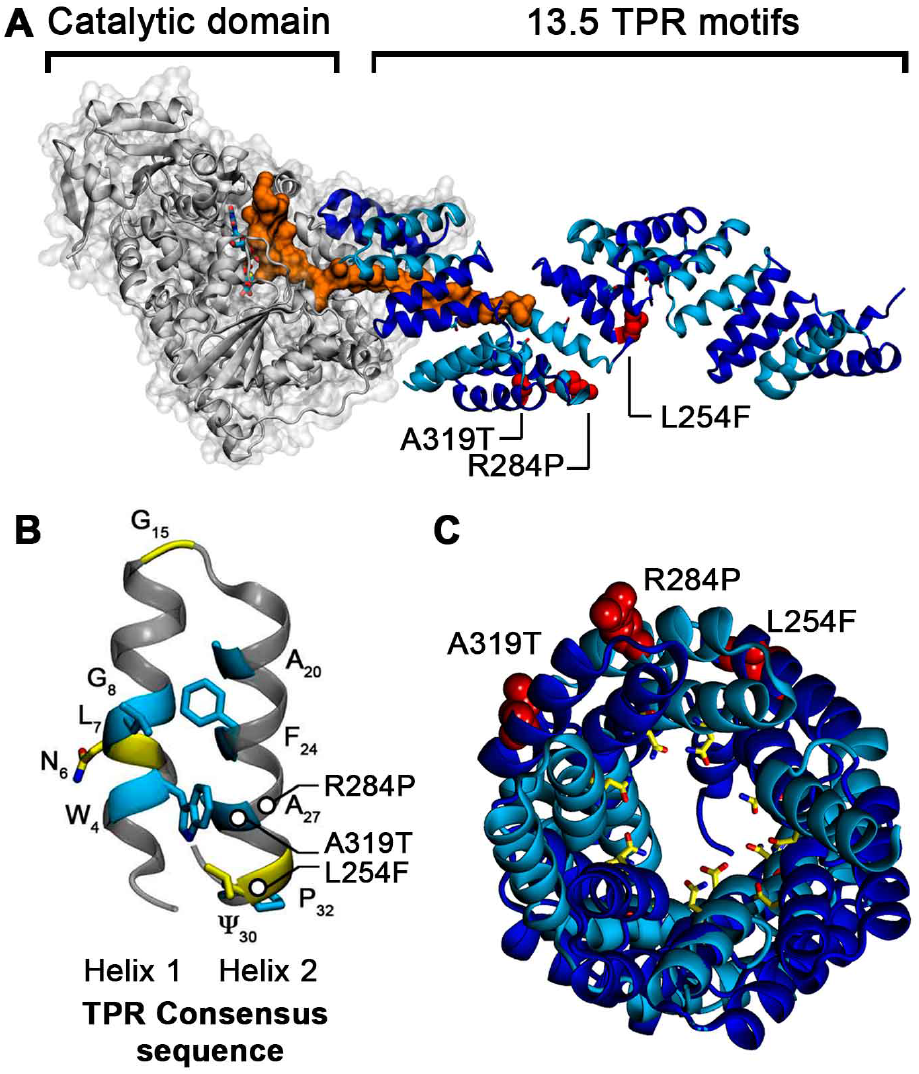
Superhelical structure of the OGT-TPR domain and location of the ID-associated mutation sites. (**A**) Structure of the OGT enzyme with the catalytic domain in grey and the TPR domain in blue cartoon representation, respectively. A short peptide belonging to a substrate protein, TAB1, is shown bound to OGT in orange (PDB id: 5LVV) [27]. The ID-associated mutations L254F, R284P and A319T are shown as red spheres. (**B**) TPR consensus sequence (CS) with the most conserved residues shown as blue sticks. The OGT enzyme contains additional conserved residues in its CS, shown as yellow sticks. (**C**) The central tunnel formed by the TPR superhelix, which is thought to form the major interaction surface for substrate proteins. The conserved N6 asparagine residues of the OGT-TPR domain, which are the main substrate interaction sites, are shown in yellow; the locations of the single point mutations are shown in red.

Here, we were interested to investigate both the global domain flexibility of the wt OGT-TPR domain as well as the effects of the ID-related single point mutations. We therefore conducted microsecond all-atom molecular dynamics simulations, both unbiased and steered, of wt and mutant OGT proteins and analysed the effect of the mutations on the dynamic properties of the domain. Our simulations first establish the TPR domain as a reversibly elastic nanospring. Furthermore, our results show that each of the single mutations alters the conformational dynamics of the domain in a different way, and leads to distinct changes in the overall biomechanical properties of OGT-TPR, while all of them display a strong divergence from the wt. These modified dynamics may play an important role in the capacity of the OGT enzyme to bind its various substrate proteins and therefore help to explain the ID phenotype of these mutants. Moreover, our findings may provide information how engineered TPR proteins could be conferred with fine-tuned dynamic properties during their structural design.

## RESULTS & DISCUSSION

### The OGT-TPR domain is a protein nanospring with fully reversible elasticity

To characterise the elasticity of the OGT-TPR domain and to ascertain if its folded structure remains intact upon enforced elongation, we performed steered molecular dynamics (SMD) simulations on wild-type (wt) OGT-TPR [21] and the ID-associated mutants [14]. The available OGT-TPR crystal structures include sequence positions 26-410, comprising ten complete TPR units (TPR_2-11_) and two partially resolved repeats. We used a moving harmonic potential of ~1.25 kcal mol^−1^ Å^−2^, attached to the C-terminal end of the TPR domain (TPR11), at a velocity of 1 Å ns^−1^ to increase its separation from the fixed N-terminus (TPR2) and thereby elongate the domain. We then extracted four independent extended conformations obtained from the trajectory under force and allowed the OGT-TPR domain to relax its conformation in further unbiased simulations.

As shown in Fig. 2A, all of the elongated conformations of wt OGT-TPR relax back to their original end-to-end distance on very short timescales, only spanning few ns. Although the fluctuation level around the equilibrium distance is relatively high (reflecting the high flexibility of the TPR domain), the final states regain conformations close or even identical to the original domain extension observed in the crystal structure. We thus find that the wt TPR domain shows fully reversible elasticity up to elongations of ~145% of its original length. During expansion, no rupture events occur that might affect the intramolecular contacts that are essential for maintaining its structure. This level of elasticity is similar to the reversibly elastic behaviour previously described for the HEAT repeat protein importin-β [3,28].

**Figure 2.**
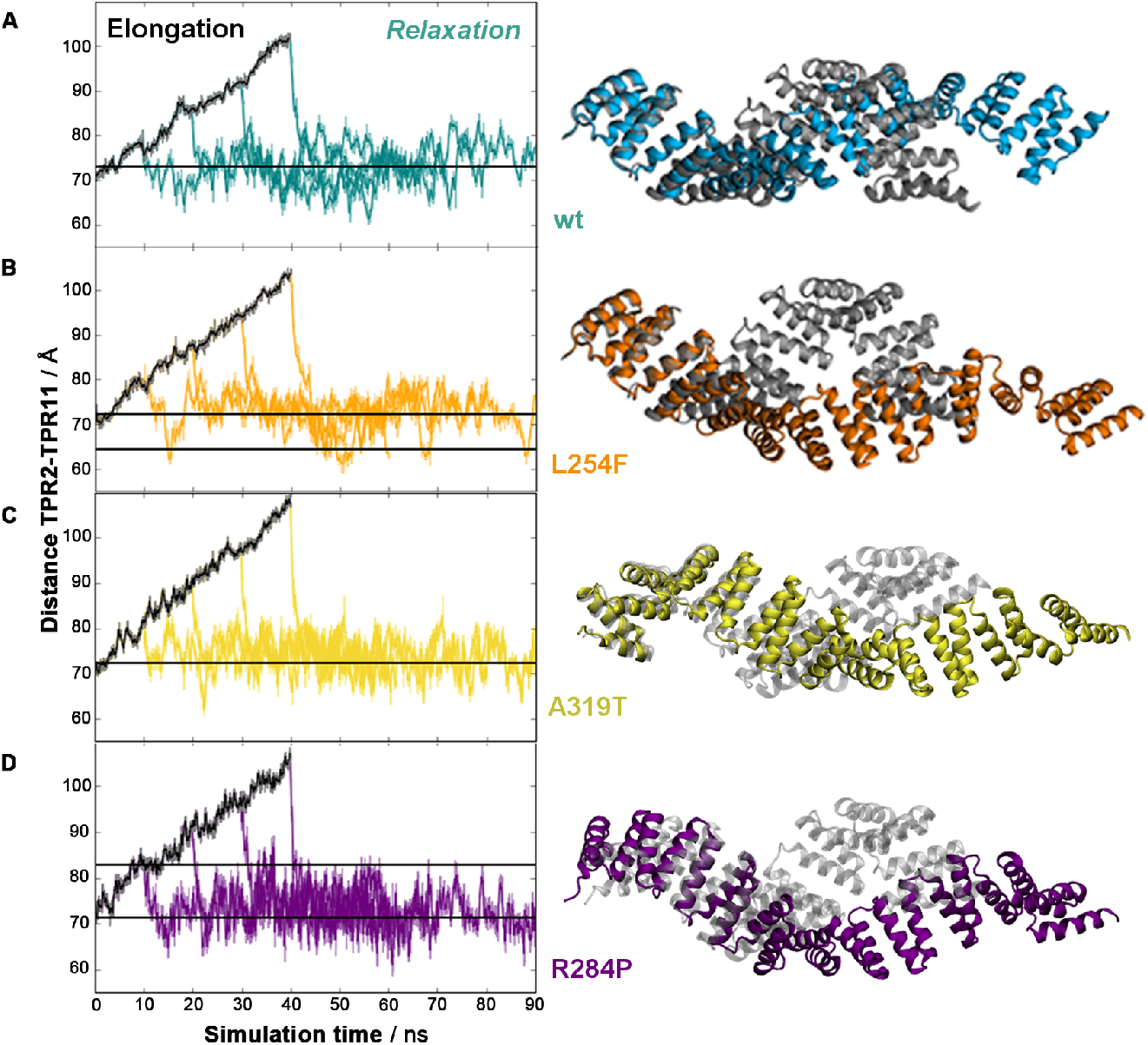
Reversible elasticity of the OGT-TPR domain. **(A, left)** Elongation of the domain under steered MD simulations (black) and subsequent relaxation of the domain back to its original length starting from four different points on the elongation trajectory (teal). **(A, right)** Maximally elongated state from which elastic relaxation to the original conformation is observed (teal) and comparison to the starting structure (grey). **(B, C, D)** Reversible elasticity and maximum elongation of the mutants L254F (orange), A319T (yellow), and R284P (purple).

The results thus show that the OGT-TPR superhelix displays a spring-like mechanical behaviour. Enforced extensions or distortions of the structure lead to a loading of this protein nanospring, by which energy is stored in the elongated conformation without disrupting its secondary structure or intramolecular contacts. Upon release of the driving force, the elongated superhelix elastically relaxes to its original ground state, thereby releasing the energy that was previously stored in the nanospring.

SMD simulations of the ID-related mutants [14,25] (Fig. 2B-D) show that these mutations do not disrupt the elastic spring behaviour of the domain. In fact, the ID variants can sustain slightly larger end-to-end extensions in the fully elastic regime before disruption of the secondary structure occurs. The maximum extensions we observe for the mutants are ~103.7 Å (L254F), 107.5 Å (A319T), and ~107.2 Å (R284P), compared to ~101.7 Å for the wt. Like the wt, all of the variants relax back to their original end-to-end length after release of the driving force. This suggests that the principal spring-like behaviour of the domain is robust against these single-point changes. We therefore conclude that the mutations do not incur a globally misfolded domain structure but rather lead to more subtle changes in the dynamical and biomechanical properties of the domain, which will be most relevant for protein-protein binding interactions. Support for this notion comes from recent experiments, in which only moderate deviations from the melting temperature of the wt domain were observed for these mutants [29].

### Biomechanical properties of the OGT-TPR domain and effect of ID-related mutations

To accurately determine the spring constant of the domain and thereby obtain the energy required for its elastic deformation, we conducted further equilibrium MD simulations. The wt and mutant OGT-TPR domains were each simulated for a total time of 2 μs, combining data from four replicates of 500 ns length. Fig. 3A shows the distribution of end-to-end (TPR2-11) domain distances observed during the simulations.

**Figure 3.**
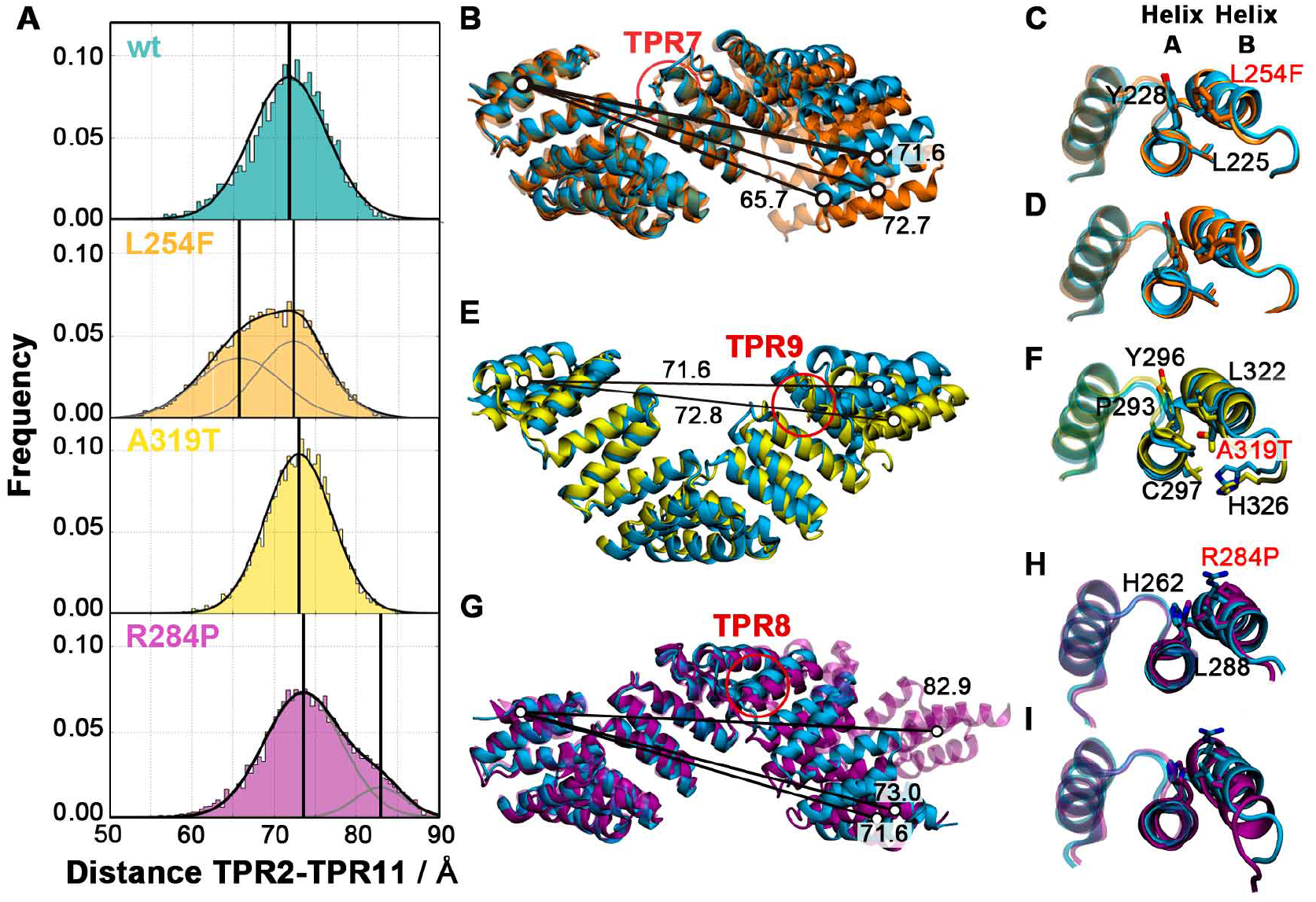
Effect of the ID-related single mutations on the elasticity of the TPR domain and local conformational changes. **(A)** End-to-end distance of the OGT-TPR domain of the wt (cyan), L254F mutant (orange), A319T mutant (yellow) and R284P mutant (purple). **(B, C, D)** Superposition of the representative equilibrium conformations of the wt (cyan) with the L254F (orange), A319T (yellow) and R284P mutant TPR-OGT domains, respectively. Indicative distances (Å) between the centres-of-mass of TPR2 and TPR11 are shown for the major mutant conformations; the mutation site is highlighted. (**E**) Representative snapshots of the two major conformations of TPR7 and the F254 sidechain compared to L254 of the wt domain. (**F**) Representative snapshot of TPR9 and the T319 sidechain compared to A319 of the wt domain. (**G**) Representative snapshots of the two major conformations of TPR8 and the P284 sidechain compared to R284 of the wt domain.

In the case of the wt, the extensions are normally distributed, reflecting the fluctuations of the TPR domain around a single equilibrium length of ~71.6 Å. Since the fluctuations of a spring are related to the spring constant by *k_spring_* = *k_B_T* / *σ*^2^, the width *σ* of the normal distribution (its standard deviation) gives rise to a spring constant of *k_WT_* = 20.80 ± 0.07 pN/nm. For comparison, the spring constant found for the HEAT repeat domain of importin-β is ~10 pN/nm [3], while that of the armadillo-repeat domain of importin-a lies between 80-120 pN/nm [4]. The spring constant of OGT-TPR signifies that an extension or compression of the OGT-TPR domain by 1 nm requires an energy input of ~6 kJ/mol, while an energy of ~20 kJ/mol is necessary to obtain the maximum elastic extension we observe in our steered simulations. Upon binding and accommodating substrate proteins of different size, the energy for the distortion of the superhelix is likely provided by the binding energy of the substrate to the TPR domain.

Importantly, this elasticity provides the domain with the ability to transiently store part of this binding energy in the form of distorting the superhelix and release this energy upon substrate dissociation. In HEAT repeat proteins, the capacity to store binding energy has been identified as a crucial factor that aids in accelerating the disassembly of protein-protein complexes. The extended surface of α-solenoid domains optimises binding selectivity by providing a multitude of specific binding interactions, whose sum however also leads to very large protein-protein binding energies [30,31]. Protein complexes of such high affinity would show exceptionally slow off-rates upon disassembly, unless some of the substrate binding energy could be stored in the deformation of the α-solenoid superhelix. In this way, extended α-solenoid domains are likely to fine-tune their protein-protein binding thermodynamics [30–32].

The distributions of end-to-end distances of the ID-related domain variants are displayed in Fig. 3A. In contrast to the wt, the L254F variant shows a partition into two populations with different average extension. The main population has an average extension of ~72.7 Å, while a second Gaussian distribution is observed around ~65.7 Å. The end-to-end distances of the R284P mutant domain also display a separation into two populations. Here, the main population has an extension similar to the wt (~73.0 Å), while the secondary population shows an increased length of ~82.9 Å. The end-to-end distance distribution of the A319T variant does not display a separation into sub-populations, and its mean remains near the wt value (~72.8 Å). However, the width of the normal distribution is markedly reduced compared to wt, indicating a modification of its nanospring behaviour. We therefore derived the spring constants for all the major conformational populations of the mutants.

The major species of the L254F mutant has a spring constant similar to the wt (*k_LF1_* = 21.99 ± 0.14 pN/nm) while the shorter population shows a markedly softened spring constant of *k_LF2_* = 12.61 ± 0.08 pN/nm. For the R284P mutant, the lengthened population also reflects a softer nanospring with a spring constant of *k_RP2_* = 17.02 ± 0.13 pN/nm, while the major population remains close to the wild-type (*k_RP1_* = 19.92 ± 0.01 pN/nm). By contrast, the narrower distribution of lengths observed for the A319T variant emerges due to a rigidified nanospring (*k_AT_* = 24.90 ± 0.11 pN/nm).

These results show that all of the ID-related single point mutations in the TPR domain induce substantial changes in the biomechanical properties of the domain. The malleability of the domain, and its capacity to adapt to different substrate proteins, is key to enabling the function of the OGT enzyme, however. Furthermore, as shown previously for HEAT repeat proteins [30,31], the nanospring character of α-solenoid domains is crucial for the reversibility of protein-protein binding during substrate release by enabling the transient storage and release of binding energy. Our finding that all of the ID-related OGT-TPR mutants exhibit a significant alteration in their spring-like behaviour thus indicates likely defects in their capacity to bind and efficiently release substrates.

### Local conformational effects propagate into globally altered L254F and R284P states

While all the mutations lead to a substantial modification of the spring constant of the OGT-TPR domain, two mutants, L254F and R284P, additionally show populations that deviate from the overall wt equilibrium length. We were therefore interested how these single-point mutations propagate into a global conformational change of the domain. We monitored the local geometry around the mutated site using three structural determinants of the individual repeats (see Fig. S2 for a graphical representation): the intra-TPR distance (distance between the Cα atoms of TPR unit positions Ψ1 and Ψ30), the inter-TPR distance (distance between the centres of mass of consecutive repeats), and the angle formed by the Cα atoms of position Ψ_30_ of the previous repeat and the positions Ψ_1_ and Ψ_30_ of the mutated TPR repeat (B-A’-B’ angle, Fig. S2) [14]. This angle quantifies the turn between repeats, which contributes to the formation of the global TPR superhelix. As measures of the global domain conformation, we used the end-to-end distance of the domain, as before, as well as its root mean square deviation (RMSD) during the simulations.

In the wt domain, TPR7 shows an intra-TPR7 distance of ~6.63 ± 0.39 Å and a 6B-7A-7B angle of 107.80° ± 4.20°. The L254 side chain is buried between TPR7 helices A and B, establishing van der Waals interactions with the side chains of L225 and Y228. Its first side chain dihedral angle, X^1^, adopts a single conformation at −72.24° ± 12.73°. By contrast, the bulkier Phe side chain in the L254F mutant can adopt three conformations around this dihedral angle – two major orientations (with χ_1_ = −54.51° ± 13.94°, termed LF1, and χ_1_ = 69.58° ± 9.74°, termed LF2, shown in Fig. 3E) as well as a transient state (χ_1_ = −167.53° ± 12.19°, LF3), as previously reported in Gundogdu et al. [14] (Fig. 4A). In both the wt and L254F crystal structures, only the LF1 conformation is seen. In the mutant LF1 conformation, the phenyl moiety interacts with the side chains of L225, Y228 and R245. The wt B6-A7-B7 angle is maintained (113.53° ± 5.51°), while the intra-TPR7 distance (6.83 ± 0.50 Å) remains close to the wt (Figs. 4A, S9). The LF2 conformation of the mutant shows an increase in the intra-TPR7 distance (to 8.83 ± 0.38 Å) and a reduced B6-A7-B7 angle (93.88° ± 4.91°). These local conformational changes enable the phenyl moiety of the mutant to wedge in between the TPR7 helices and interact with the side chains of N224, L225 and Y228 (Fig. 3E). The two major different local conformational states within mutant repeat TPR7 propagate to the neighbouring repeat modules and, as a consequence, modify the overall geometry of the domain. The global end-to-end distance distribution of the L254F mutant is thus bimodal, with two Gaussians reflecting the two major conformations of the F254 residue (Figs. 4 and 3A). In the case of the A319T mutation, we find that the rigidification of the nanospring is due to the formation of an additional hydrogen bond between the side chain of T319 on TPR9 helix B and the backbone of Y296 on TPR9 helix A (Fig. 3F).

**Figure 4.**
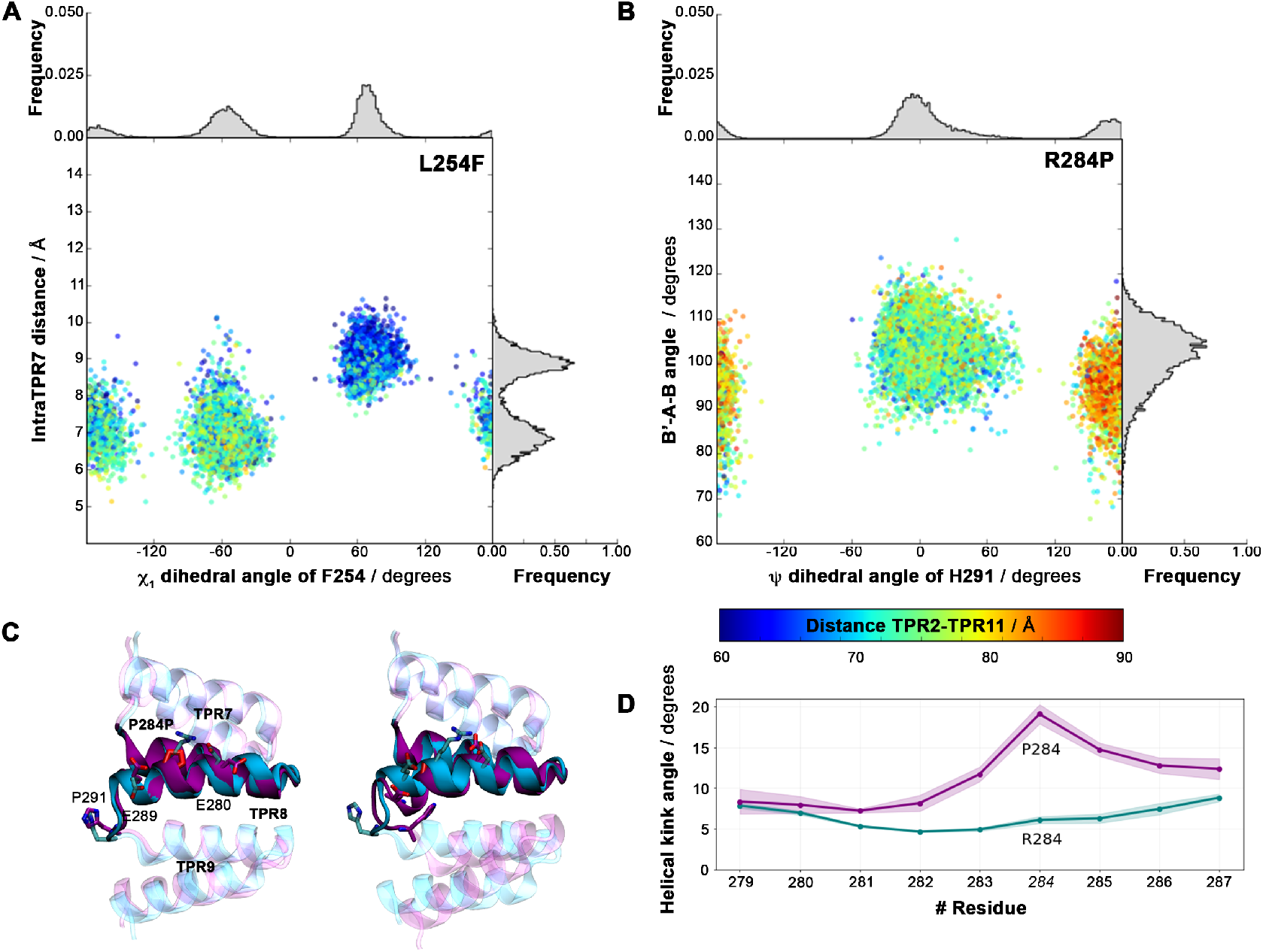
Local conformational populations of the L254F and R284 mutants. **(A, B)** Major local structural determinants are shown on the x- and y-axis, the global TPR2–TPR11 distance is colour-coded. A clear correlation between the local conformation and the global length is seen. **(A)** The x-axis displays the X1 dihedral angle of residue F254, the y-axis the intra-TPR7 distance; normalised histograms in grey show the relative distributions of the local populations. **(B)** The ψ dihedral angle of residue H291 is shown on the x-axis, the angle between consensus position 30 of TPR8 and positions 2 and 30 of TPR9 (B’-A-B) on the y-axis; normalised histograms display the relative distributions of the populations. **(C)** Representative conformations of the two populations found for R284P. The R284P and wt domains are shown in purple and cyan cartoon and sticks, respectively. **(C)** Kink angle between successive residues in TPR8 helix B for R284P (purple) and wt (cyan).

The R284P mutation, located in repeat 8 (TPR8) at position X_26_ outside the TPR consensus sequence, introduces a proline residue in the middle of TPR helix B. This mutation restricts the mobility of the P284 sidechain and abolishes the salt bridge between R284 and residues E280 and E289 from the same helix. Additionally, a proline residue cannot establish the wt hydrogen bond with the previous helix turn, distorting the helical domain. This increases the distance between the backbone O atom of E280 and the N atom of the P284 side chain from 3.00 ± 0.16 Å in the wt to 4.62 ± 0.25 Å in the mutant. Additionally, in a minor population, the helix develops a kink of 19.1 ± 1.0 degrees (Fig. 3G, 4D). The altered geometry of TPR8 influences the neighbouring TPR unit by changing the inter-repeat angle and modifying the conformation of residue H291 on TPR9 (Fig. 4B). In the R284P minor conformation, the H291 side chain resides between the side chain of F292 and the backbone of the A285, while it is solvent-exposed in both the wt and the major conformation of R284P (Fig. 4C). The substantial local rearrangements in the smaller population of R284P are then propagated into a globally elongated domain conformation.

## CONCLUSION

TPR domains are involved in many key biological processes through their ability to bind selectively to an array of different protein partners. The stacking of repeat units, forming a superhelical global structure, provides TPR domains with high flexibility while retaining a robust protein fold. Here, we have characterised the nanospring character of the OGT-TPR domain and found it to show fully reversible elasticity over a wide range of domain extensions. A small number of single-point mutations within this domain are associated with ID phenotypes. Interestingly, while not all of these mutations lead to changes in the equilibrium domain conformation, they all show strong deviations from the wild-type elasticity and dynamics. These differences also impact on the energetics of the global conformational changes of the domain that underpin substrate binding and release. Some of these effects may not be detectable in crystal structures, since the average, or dominant, domain conformation often remains unchanged. Taken together, our results suggest that the mutations are likely to display substantial defects upon substrate interaction, due to their altered flexibility and conformational energetics. Our findings may provide a clue towards the ID phenotype of these single-point mutations, which are all distal from the OGT active site but locate to an important part of its substrate binding domain. In addition, they could bring a new perspective to the design of engineered TPR proteins [18,33] by showing how the dynamics of the domains can be fine-tuned through the introduction of subtle modifications into the repeat sequences.

## METHODS

### System Setup

A shortened construct (residues 22 – 413 from the OGT protein) was chosen to model the wt and mutant OGT-TPR domains. This model protein includes repeats TPR2 to TPR11. The constructs were capped and solvated in a triclinic box of 116.6 × 116.6 × 116.6 Å. Na^+^ and Cl^−^ ions were added to neutralise the system and reach a physiological concentration of 0.15 mM NaCl. The amber99sb-ildn force field [34] and virtual sites for hydrogen atoms [35] were used for the protein. The TIP3P water model was used to model the solvent molecules [36] and Joung and Cheatham III parameters were used to model the ions [37].

### Unbiased Molecular Dynamics Simulations

Molecular simulations were performed with the GROMACS molecular dynamics package version 5.1.5 [38]. For each system, the geometry was minimized in four cycles that combined 3500 steps of steepest descent algorithm followed by 4500 of conjugate gradient. Thermalization of the system was performed in 10 steps of 2 ns, where the temperature was gradually increased from 50 K to 298 K, while the protein was restrained with a force constant of 10 kJ mol^−1^ Å^−2^. Production runs consisted of four replicates of simulations of 500 ns length for each system (accounting for a total of 2.0 μs of simulation time), using an integration time-step of 4 fs.

The temperature was kept constant by weakly coupling (t = 0.1 ps) the protein and solvent separately to a temperature bath of 298 K with the velocity-rescale thermostat of Bussi et al. [39]. The pressure was kept constant at 1 bar using isotropic Berendsen coupling [40]. Long-range electrostatic interactions were calculated using the smooth particle mesh Ewald method [41] beyond a short-range Coulomb cut-off of 10 Å. A 10 Å cut-off was also employed for Lennard-Jones interactions. The LINCS algorithm [42] was used to restrain the bonds involving hydrogen and SETTLE algorithm [43] was used to constrain bond lengths and angles of water molecules. Periodic boundary conditions were applied.

### Analysis of the trajectories and estimation of the spring force constant

We used MDAnalysis [44,45] and MDtraj [46] to analyse the trajectories. To estimate the spring force constant, we used the protocol described in Kappel et al. [3]. Errors bars were obtained by bootstrap analysis of the mean of the widths of the Gaussian distributions (1000 cycles).

### TPR domain elastic behaviour

A slightly different system setup was employed for our Steered Molecular Dynamics simulations [47]. To ensure that any extended state of the proteins fits into the simulation box, proteins were oriented along the z-axis of box vectors. The simulation box was subsequently extended by 60 Å along the z-axis, resulting in a box of 85.5 × 85 × 180 Å. Afterwards, the systems were solvated and a physiological concentration of Na^+^ and Cl^−^ ions was added, replicating the protocols used for the unbiased molecular dynamics simulations. The aforementioned thermalisation and equilibration protocols were also used here. The stretching protocol consisted in fixing the C atoms of the helix TPR1B (N-terminal) using a force constant of 10 kJmol^−1^Å^−2^ and applying a pulling potential of 5 kJmol^−1^Å^−2^ to displace helix TPR12A (C-terminal) at a constant velocity of 1 Å ns^−1^ in the *z*-direction.

## ACKNOWLEDGMENTS

We thank Mehmet Gundogdu, Caroline Fässler, Geoff Barton and Daan van Aalten for fruitful discussions as well as Neil Thomson for critically reading the manuscript. We gratefully acknowledge funding from the Wellcome Trust (ISSF award WT097818MF) and from SUPA (Scottish Universities’ Physics Alliance).

