## Supplementary Information for "Disease-related single-point mutations alter the global dynamics of a tetratricopeptide (TPR) a-solenoid domain"

Index

Figure S1. Alignment of the OGT-TPR repeat sequence	S2

Figure S2. Geometrical descriptors of the TPR domains used in this publication.	S2

Figure S3. Time evolution of the RMSD of the backbone of the protein.	S3

Figure S4. Time evolution of the TPR2 – TPR11 distance for the four studied proteins.	S3

Figure S5. Geometrical parameters of the wt OGT-TPR domains.	S4

Figure S6. Geometrical parameters of the L254F OGT-TPR domains.	S5

Figure S7. Geometrical parameters of the L254F OGT-TPR domains.	S6

Figure S8. Geometrical parameters of the L254F OGT-TPR domains.	S7

Figure S9. Distortion caused by the single point mutation L254F.	S8

Figure S10. Distortion caused by the single point mutation R284P.	S9

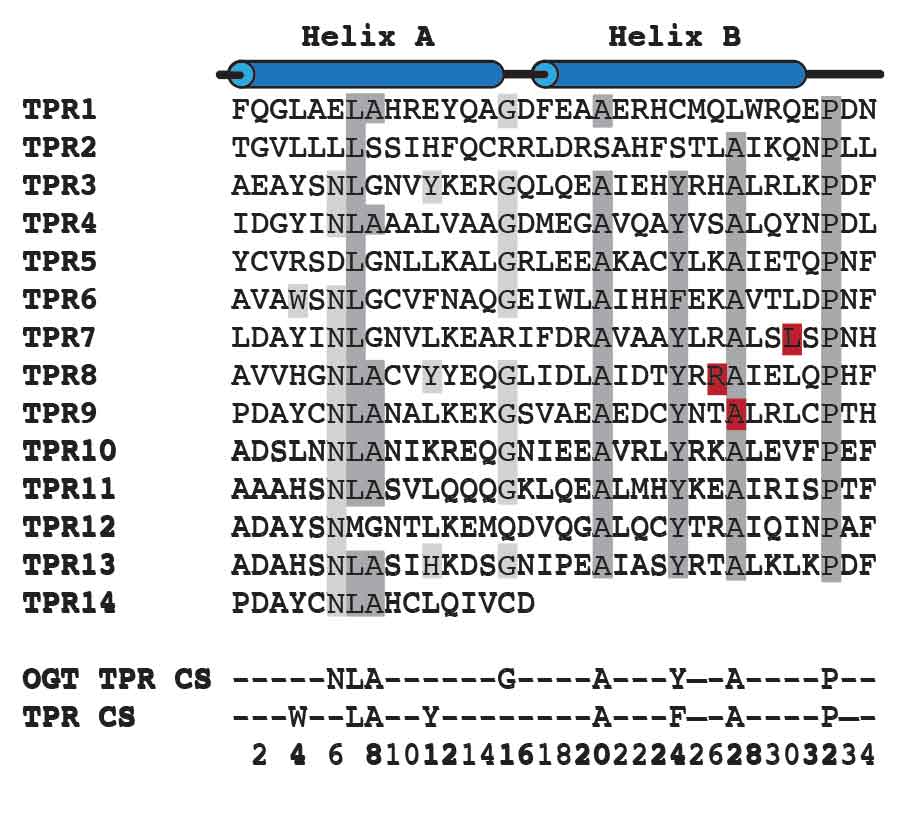

**Figure S1.** **Alignment of OGT-TPR sequence**. The single point mutations are highlighted in red.

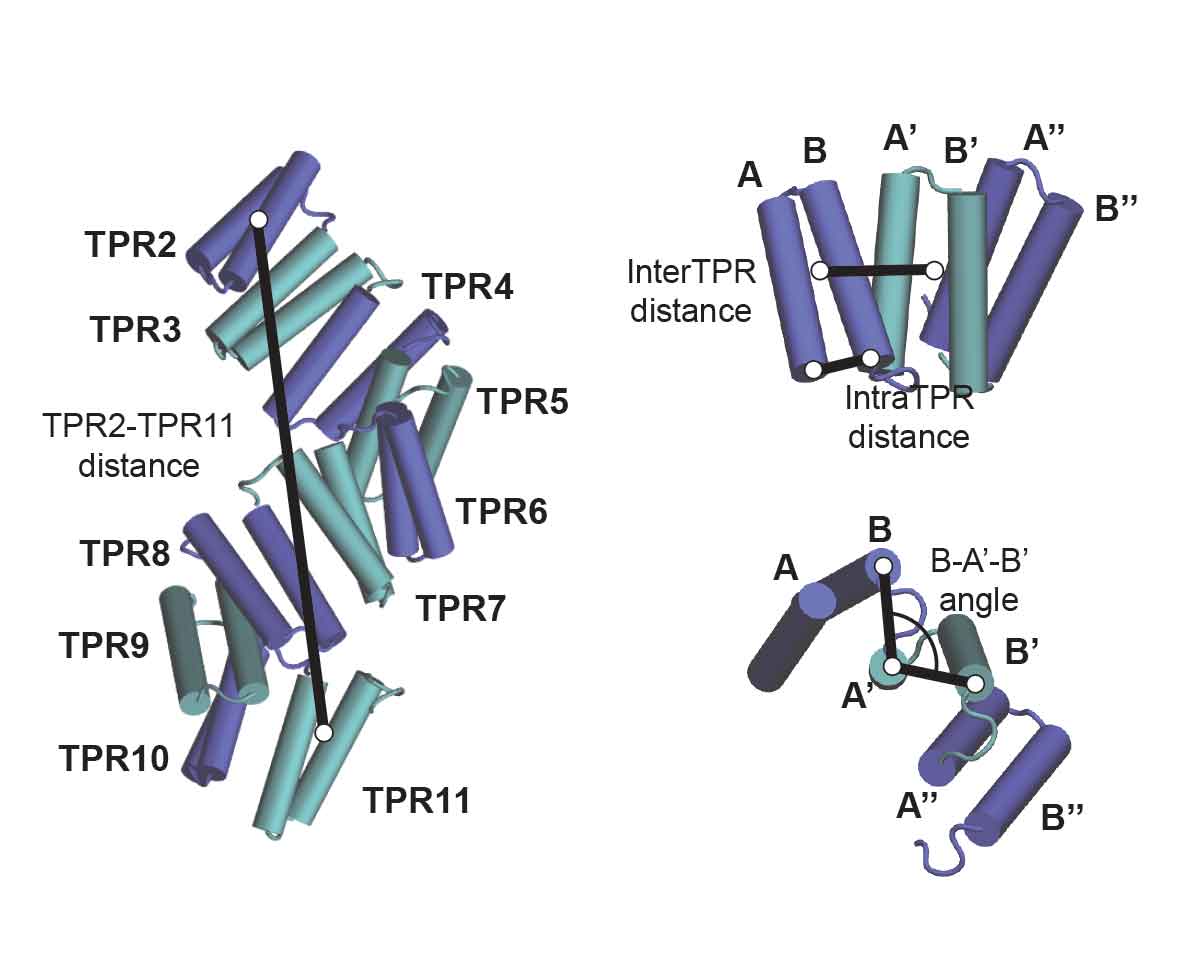

**Figure S2. Geometrical descriptors of the TPR domains**. The intra-TPR distance is calculated as the distance between the Cα atoms of the positions 0 and 30 of the TPR sequence of each domain. The inter-TPR distance is calculated as the distance between the centres-of-mass (COM) of adjacent TPR domains. The B’-A-B angle is the angle formed by the positions of the positions 0 and 30 of the TPR sequence of each domain and the position 30 of the previous repeat. The TPR2-TPR11 distance is defined by the distance between the COM of the repeats TRP2 and TPR11.

| **Wild type**  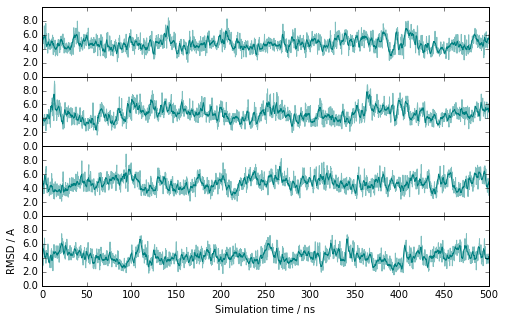 | **L254F mutant**  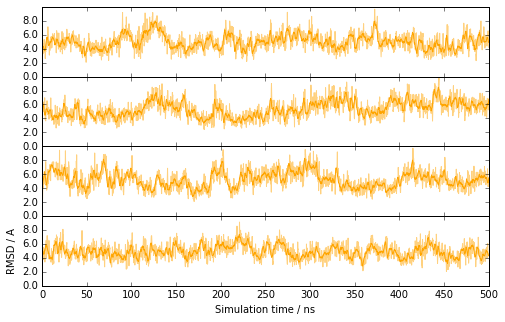 |
| --- | --- |
| **A319T mutant**  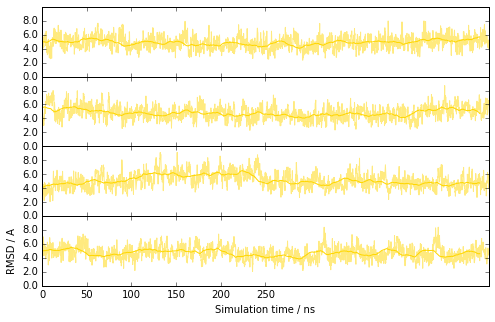 | **R284P mutant**  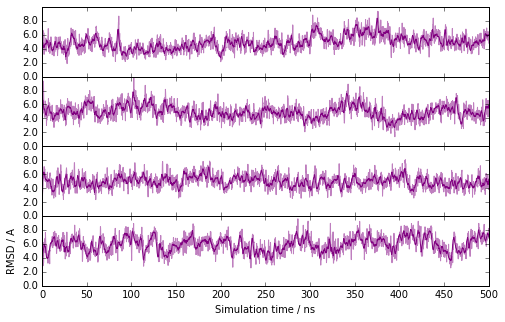 |

**Figure S3. Time evolution of the RMSD of the backbone of the protein**. Wild type, L254F, A319T and R284P domains are shown in cyan, orange, yellow and purple plots respectively.

| **Wild type**  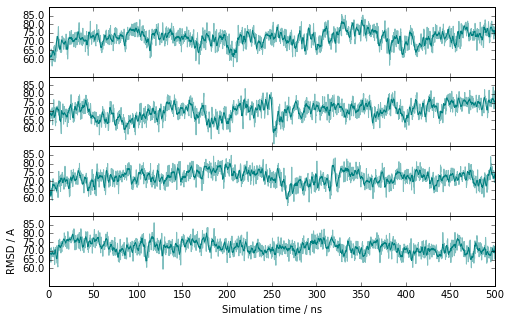 | **L254F mutant**  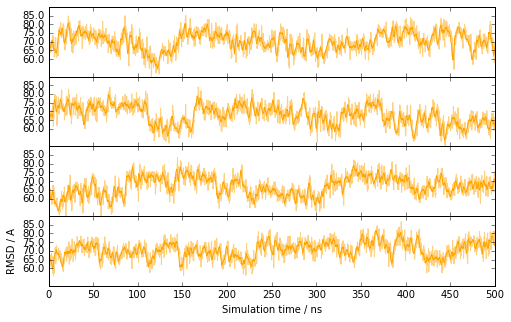 |
| --- | --- |
| **A319T mutant**  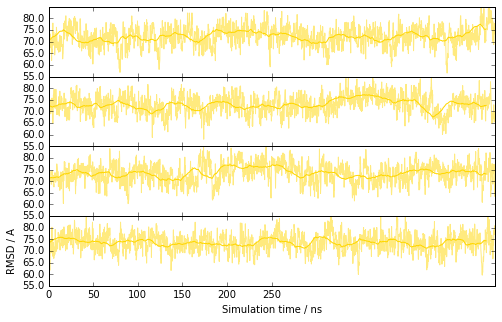 | **R284P mutant**  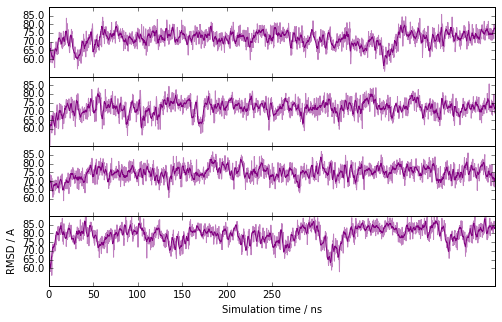 |

**Figure S4. Time evolution of the TPR2 – TPR11 distance for the four studied proteins.** Wild type, L254F, A319T and R284P domains are shown in cyan, orange, yellow and purple plots respectively.

**IntraTPR Distance**

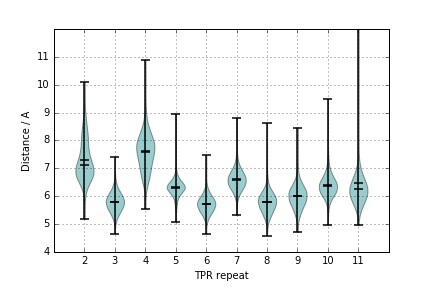

**InterTPR Distance**

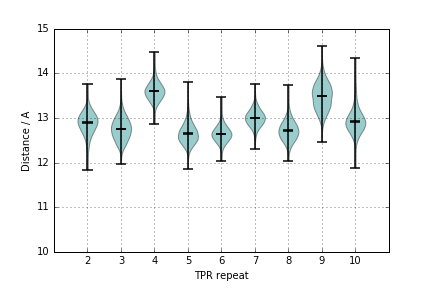

**B-A’-B’ Angle**

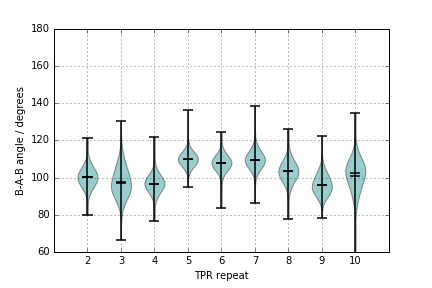

**Figure S5. Geometrical parameters of the *wt* OGT-TPR domains.** Violin plots highlight the distributions of each repeat and the mean and the median values.

**IntraTPR Distance**

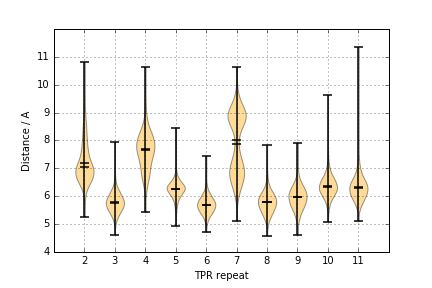

**InterTPR Distance**

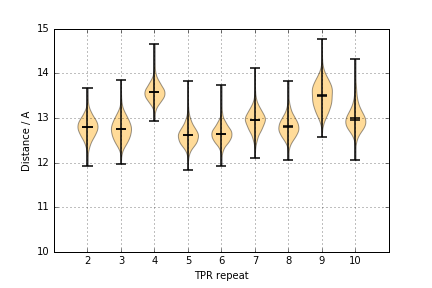

**B-A’-B’ Angle**

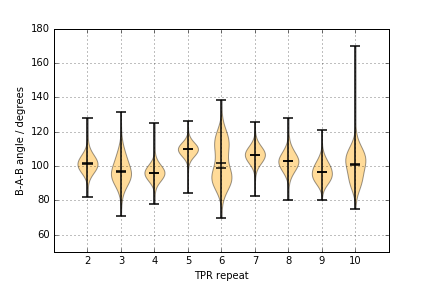

**Figure S6. Geometrical parameters of the *L254F* OGT-TPR domains.** Violin plots highlight the distributions of each repeat and the mean and the median values.

**IntraTPR Distance**

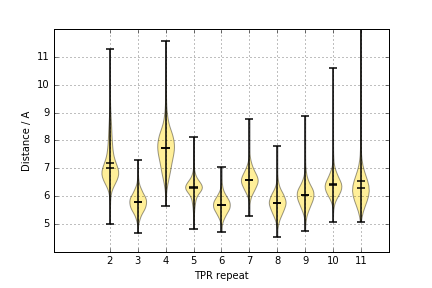

**InterTPR Distance**

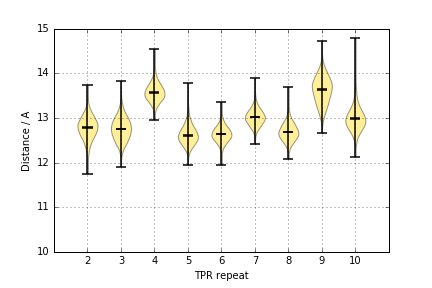

**B-A’-B’ Angle**

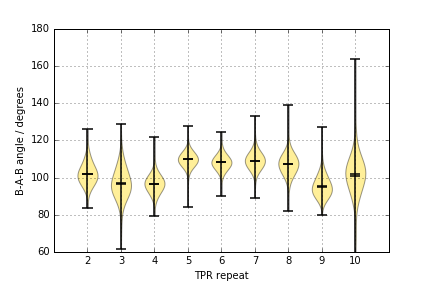

**Figure S7. Geometrical parameters of the *A319T* OGT-TPR domains.** Violin plots highlight the distributions of each repeat and the mean and the median values.

**IntraTPR Distance**

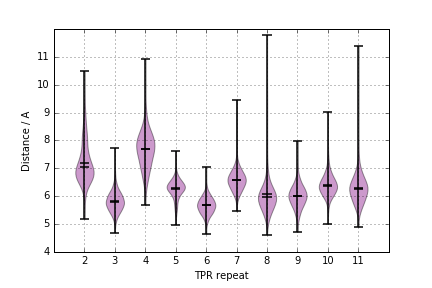

**InterTPR Distance**

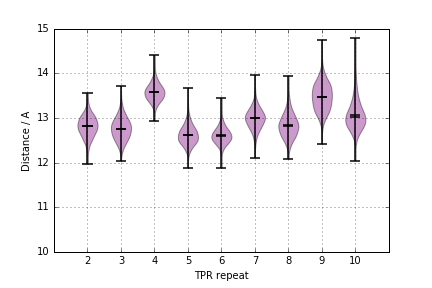

**B-A’-B’ Angle**

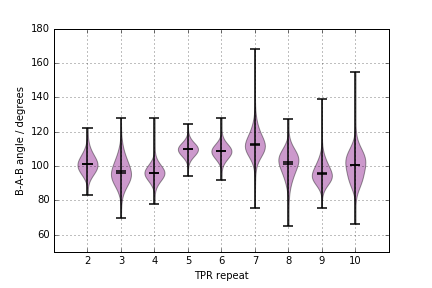

**Figure S8. Geometrical parameters of the *R284P* OGT-TPR domains**. Violin plots highlight the distributions of each repeat and the mean and the median values.

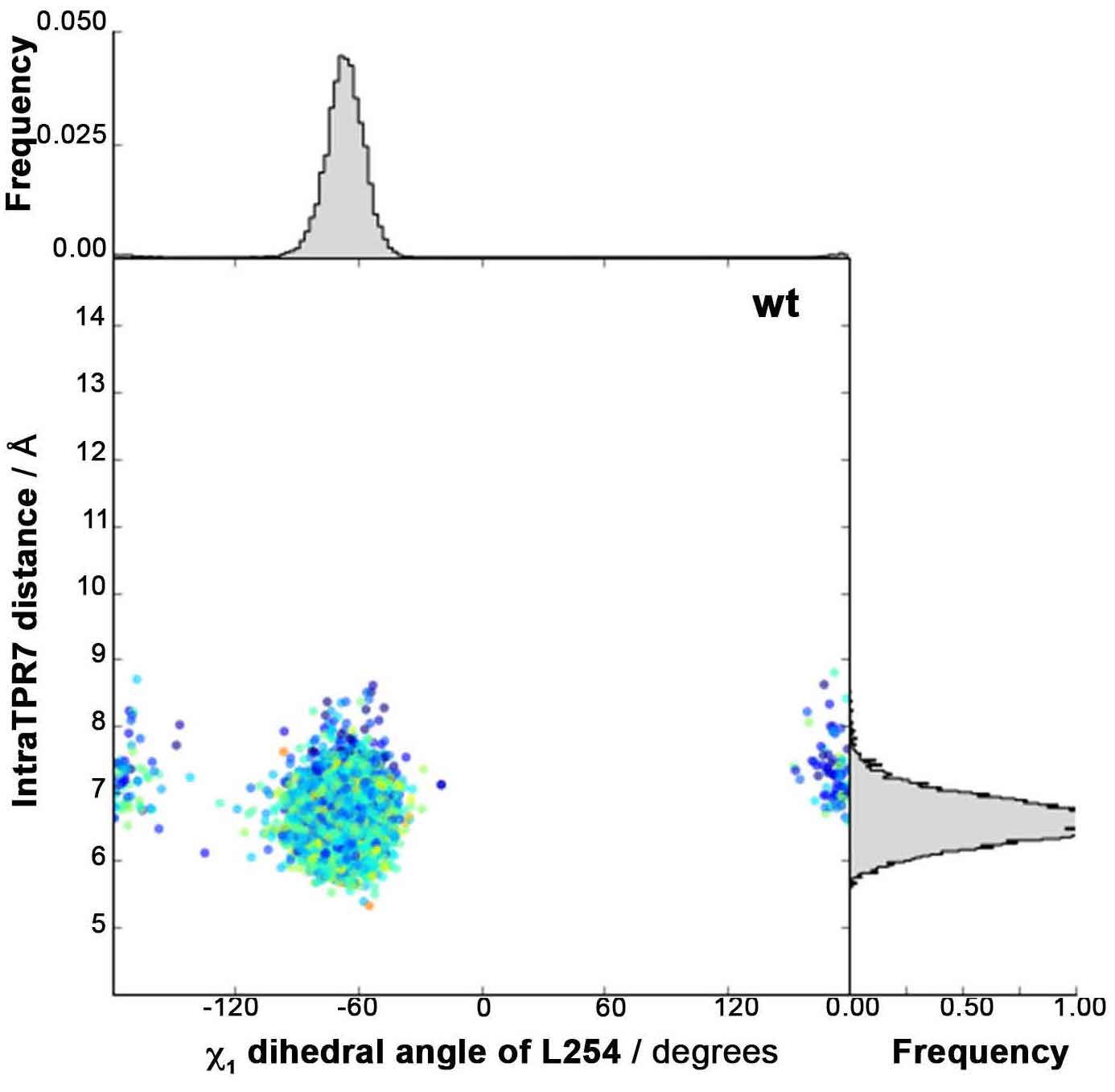

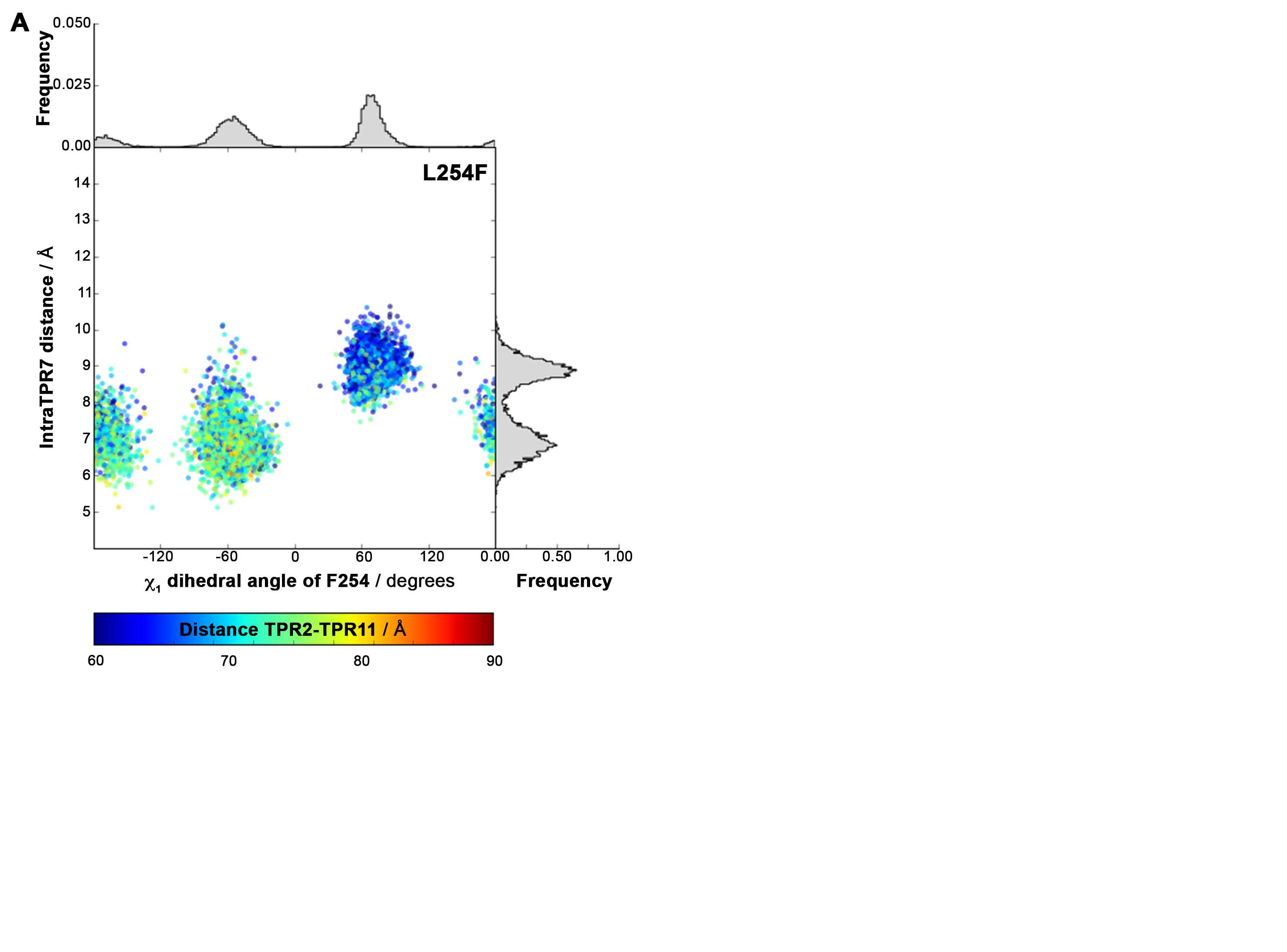

**Figure S9.** Distortion caused by the single point mutation L254F on the Chi-1 dihedral angle of the residue 254, the intra-TPR7 distance and the TPR2-TPR11 distance.

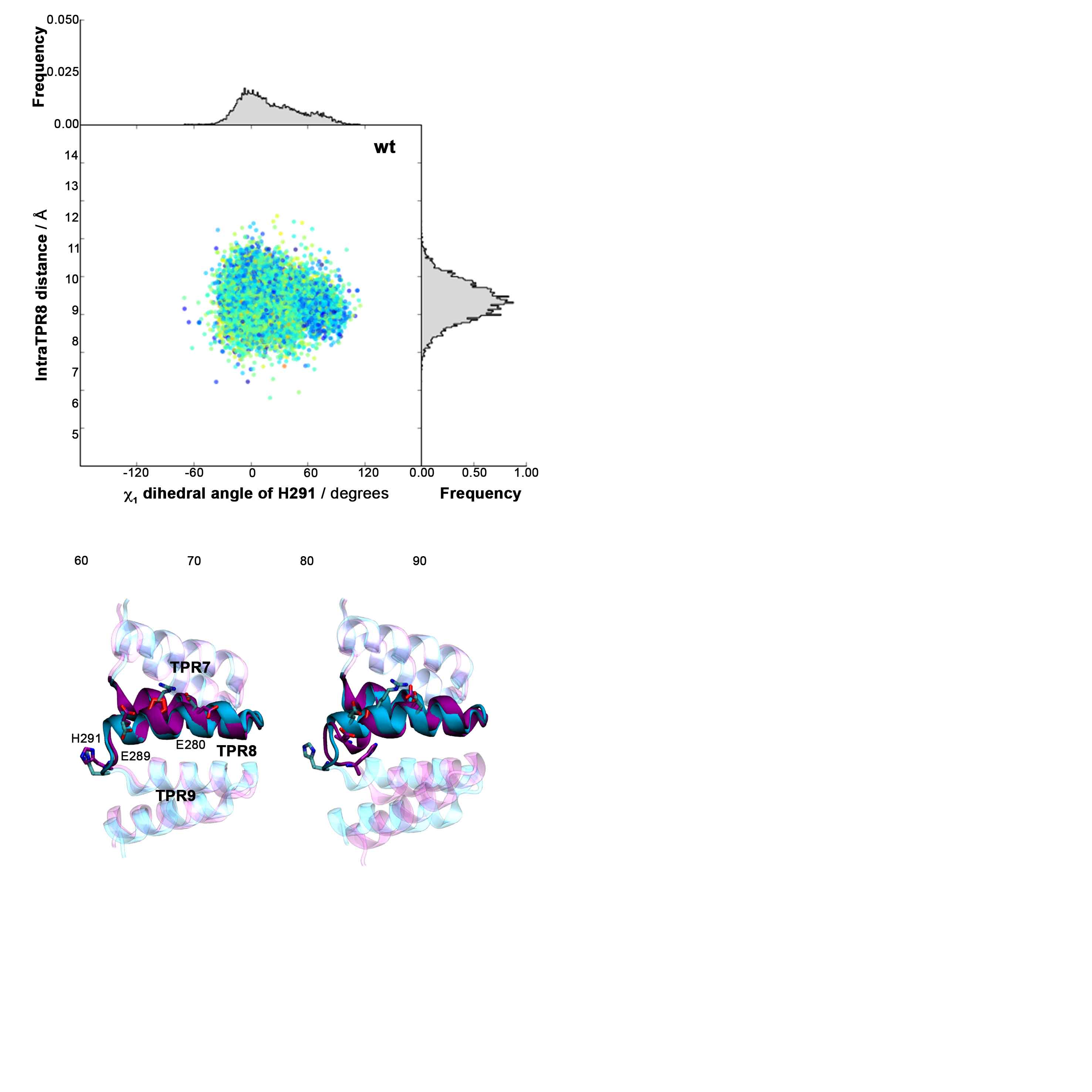

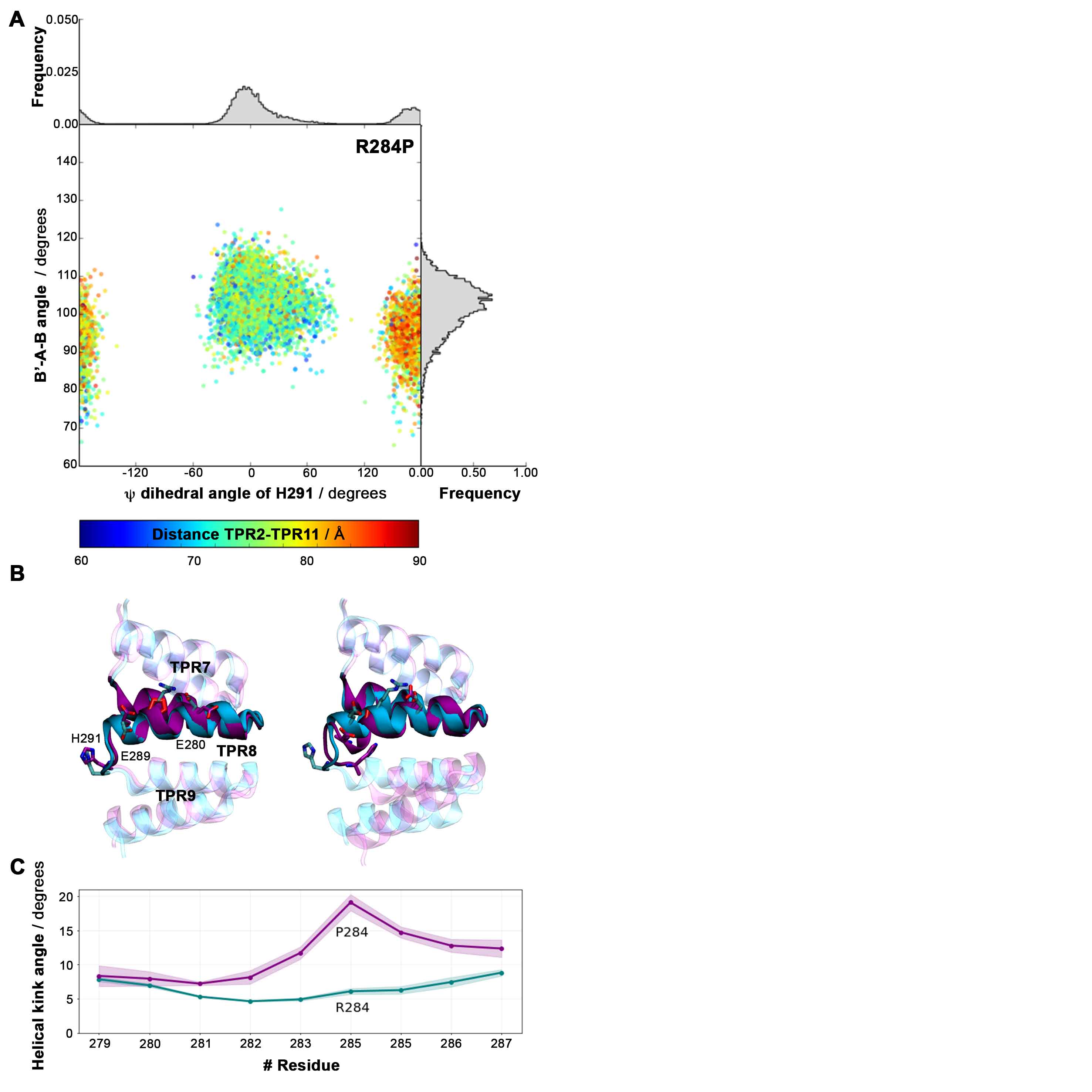

**Figure S10.** Distortion caused by the single point mutation R284P on the Chi-1 dihedral angle of the residue 291, the B’-A-B angle between TPR8 and TPR9 and the TPR2-TPR11 distance.
